# Slow conformational changes of blue light sensor BLUF proteins in milliseconds

**DOI:** 10.1101/2021.12.13.472511

**Authors:** Shunrou Tokonami, Morihiko Onose, Yusuke Nakasone, Masahide Terazima

## Abstract

BLUF (blue light sensor using flavin) proteins consist of flavin-binding BLUF domains and functional domains. Upon blue light excitation, the hydrogen bond network around the flavin chromophore changes, and the absorption spectrum in the visible region exhibits red-shift. Ultimately, the light information received in the BLUF domain is transmitted to the functional region. It has been believed that this red-shift is complete within nanoseconds. Contrary to this commonly accepted scheme, in this study, slow reaction kinetics were discovered in milliseconds (τ_1_- and τ_2_-phase) for all the BLUF proteins examined (AppA, OaPAC, BlrP1, YcgF, PapB, SyPixD, and TePixD). Despite extensive reports on BLUF, this is the first clear observation of the BLUF protein absorption change with the duration in the millisecond time region. From the measurements of some domain-deleted mutants of OaPAC and two chimeric mutants of PixD proteins, it was found that the slower dynamics (τ_2_-phase) are strongly affected by the size and nature of the C-terminal region adjacent to the BLUF domain. Hence, this millisecond reaction is a significant indicator of conformational changes in the C-terminal region, which is essential for the biological functions. On the other hand, the τ_1_-phase commonly exists in all BLUF proteins, including any mutants. The origin of the slow dynamics was studied using site-specific mutants. These results clearly show the importance of Trp in the BLUF domain. Based on this, a reaction scheme for the BLUF reaction is proposed.

**Graphical Abstract:** 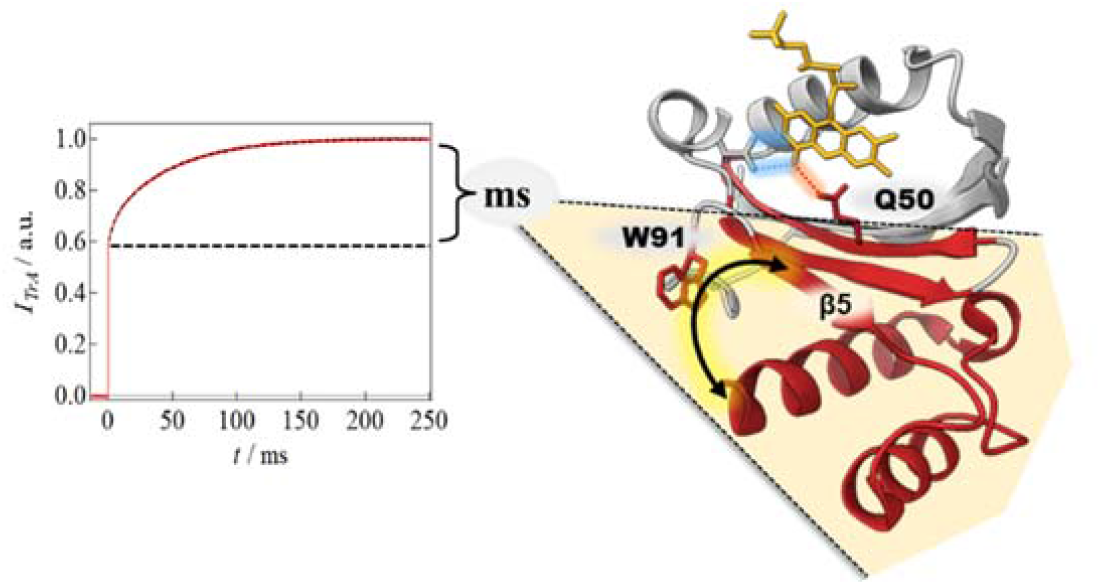

## Introduction

BLUF (blue light sensor using flavin) proteins are light sensor proteins with a BLUF domain, which consists of five β-strands, two α-helices parallel to one side of the β-sheet, and a flavin chromophore (Fig. 1(a)).^1,2^ There are two types of BLUF proteins: a short type which has no domain structure other than the BLUF domain, and a multi-domain type which has separate functional domains.^3^ The isoaloxazine ring of the flavin is bound between the two α-helices in the BLUF domain and shows hydrogen bonding to some amino acid residues, such as Asn and Gln, which are located on the α1 helix and β3 strand, respectively (e.g., N32 and Q50 in SyPixD).^4^ NMR^5^ and X-ray crystallography^6^ have shown that many BLUF proteins possess C-terminal cap helices, connecting the BLUF domain with C-terminal domains. It has been revealed that the C-terminal cap helices are important for regulating intermolecular interactions and structural changes in the C-terminal functional domains,^7,8,9^ which have roles in either enzymatic activities or the downstream transmission of light information sensed in the BLUF domain. For example, SyPixD forms a decamer with the downstream partner protein PixE, which dissociates into PixD dimers and PixE monomers upon light illumination.^10^ The interface of this complex involves a cap helix, whose conformational change is thought to be responsible for the dissociation. In OaPAC (photoactivated adenylyl cyclase from *Oscillatoria acuminata*), the BLUF domain controls the activity of the C-terminal adenylate cyclase (AC) domain via the α3 cap helix and regulates the enzymatic function of converting ATP to cAMP in a light-dependent manner.^9^ That is why the photoreactions of the BLUF proteins have attracted much attention as optogenetic tools.^11^

**Fig. 1.**
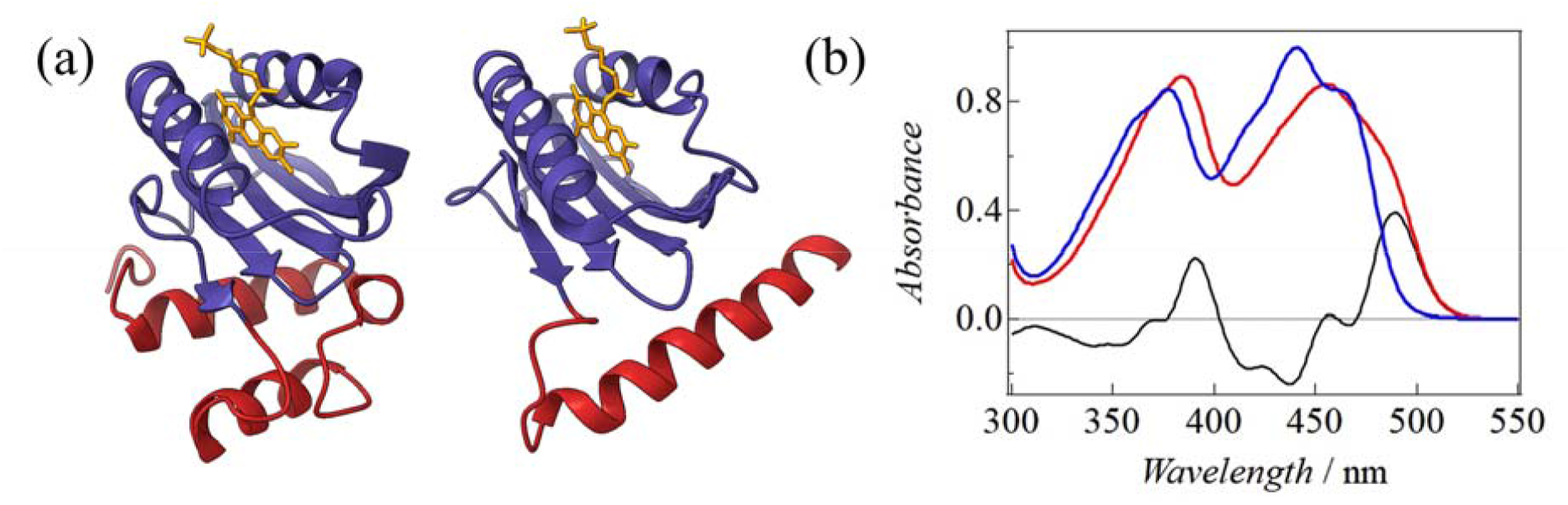
**(a)** Structures of the BLUF domains (blue); (left) SyPixD_1-150_, and (right) OaPAC_1-125_. The C-terminal cap helices are shown in red. **(b)** Typical absorption spectra of BLUF proteins (SyPixD) in the dark state (blue) and light state (red) and their difference spectra.

Because of the importance of these BLUF proteins, the reactions of the BLUF domains have been the major targets of experimental and theoretical studies.^12,13,14,15^ Although BLUF proteins are flavoproteins similar to cryptochromes and LOV (light-oxygen-voltage) proteins, the photoreaction of the chromophore flavin is very different and shows a characteristic photocycle. Upon blue light illumination, the hydrogen bond network around the flavin changes, which results in the red-shift of the absorption spectrum in the visible wavelength region.^16,17,18^ The reaction dynamics of the absorption spectrum, in particular, have been studied by transient absorption (TrA) spectroscopy.^19,20,21,22^ For example, after photoexcitation of SyPixD, proton transfer, coupled with electron transfer from the neighboring Y8 to the singlet excited state of flavin, is induced on a picosecond time scale and a flavin radical is formed.^20^ Upon returning the electrons from the radical to Y8, the orientation of the side chain of Q50 changes by rotation and/or keto-enol tautomerization,^23^ and the hydrogen bond network between the flavin and some amino acid residues of the BLUF domain is altered, resulting in the formation of a red-shifted absorption spectrum species (Fig. 1(b)).^1^ Based on previous studies, it is now widely accepted that the red-shifted state is created within nanoseconds. Indeed, the absorption changes in a slow time region, for example, milliseconds, have been reported in only two cases, and the results are controversial. A slow component, with a time constant of 5 ms for a short construct of AppA (AppA_1-156_), was once reported.^24^ However, later, an absorption change was measured in a similar time range for a slightly shorter AppA construct (AppA_5-125_), and it was reported that any slow component was not observed.^19^ Since then, there have been no further studies on the absorption changes of the BLUF proteins in the millisecond time region, and it has been common consensus that the red-shift state formation completes within nanoseconds.

Another controversial issue regarding the reaction of BLUF proteins, besides the slow absorption change, is the role of a Trp residue in the BLUF domain, which is located on β5 of the BLUF domain, near the C-terminus. It is thought to play a role in transmitting changes around the flavin to the C-terminus. For example, W104 is located close to the flavin (Trp_in_ structure) in the NMR solution structure of AppA_5-125_,^25^ whereas W104 is located away from the flavin (Trp_out_ structure) in the crystal structure of AppA_1-124_.^26^ Moreover, it has been reported that W104 shows in/out movement (flip) upon light illumination in the BLUF domain of AppA,^27^ and that this Trp is critical for biological function.^28^ On the other hand, according to the crystallographic structure of SyPixD, W91 is located close to the flavin in one of the decamer subunits, whereas, in the other subunits, W91 is located away from the flavin.^4^ These studies pointed out the importance of Trp in the BLUF reaction of PixD. However, fluorescence measurements^29^ and computational calculations^30^ indicate that the movement of this Trp is not as large as that of AppA, and that this Trp is not a crucial residue for biological function.^31^ Hence, the importance of Trp is still controversial.

In this study, we investigated slow reaction phases, monitored by light absorption in the visible wavelength region, for various BLUF proteins (AppA_1-398_, OaPAC, BlrP1, YcgF, PapB, SyPixD, and TePixD). Fig. 2(a) shows the domain structures and functions of these BLUF proteins. We report a discovery of slow reaction phases in milliseconds time range for all the BLUF proteins we studied. The slow changes are expressed by time constants of several milliseconds and/or tens of milliseconds. The millisecond component is conserved in all BLUF proteins and may be a key step in BLUF domain signaling. On the other hand, components with lifetimes of several tens of millisecond are observed in some proteins. The origin of the slow change in absorption was investigated using various mutants, and it was found that the slower dynamics reflect a conformational change at the C-terminus. Furthermore, we studied the slow component of some site-directed mutants of SyPixD, AppA_1-398_, and OaPAC, and found that the Trp residue (W91 in SyPixD, W90 in OaPAC, and W104 in AppA) is indeed involved in connecting the C-terminus to the environment of the flavin chromophore in the millisecond time region.

**Fig. 2.**
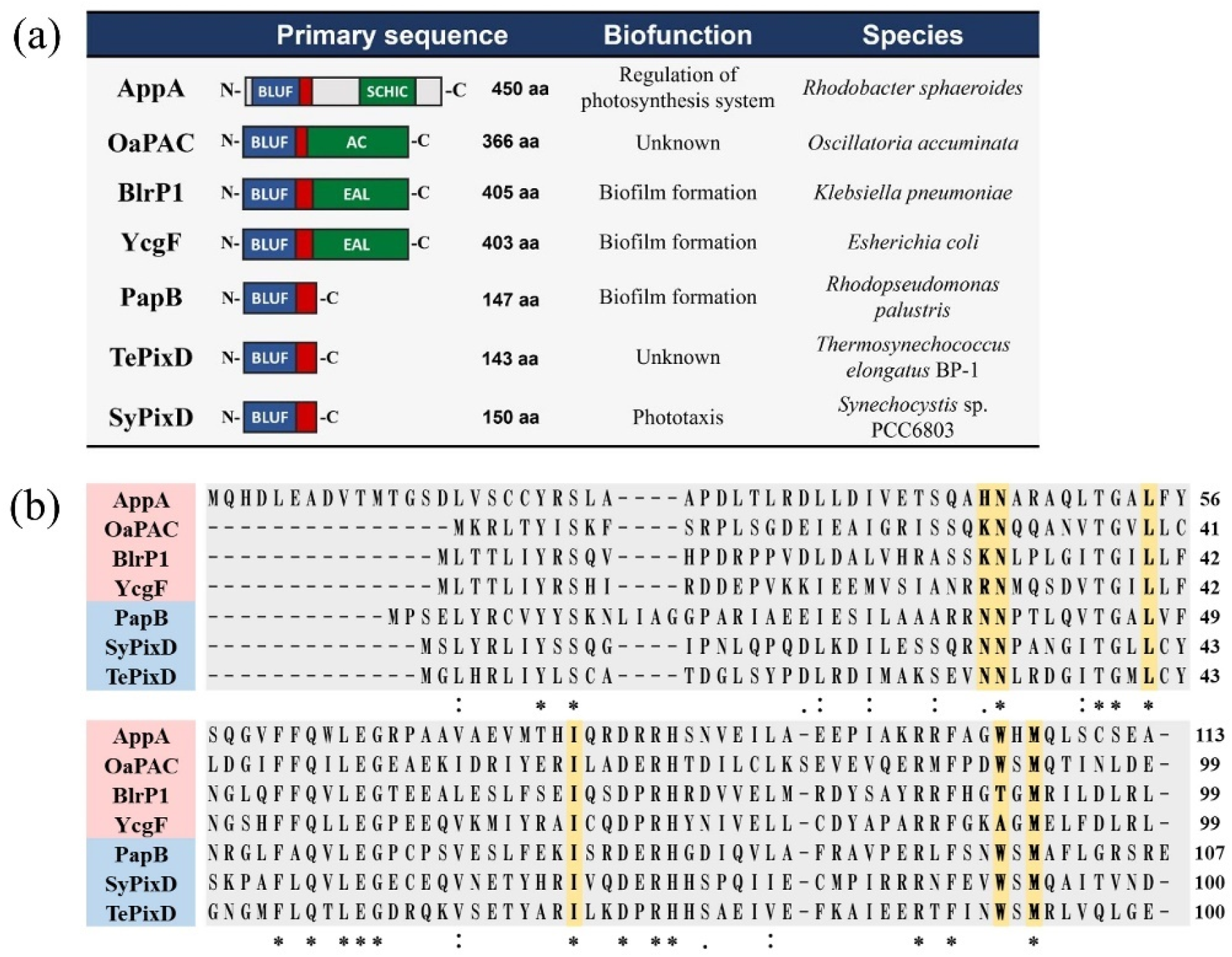
**(a)** Domain structures, biofunctions, and species of the BLUF proteins investigated in this study. **(b)** Sequence alignment of the BLUF domains (https://www.ebi.ac.uk/Tools/msa/). The positions mutated in this study are highlighted by yellow and fully conserved residues are marked by asterisks (*).

## Material and Methods

### Sample preparation

BLUF proteins used in this study were expressed in *E. coli* and purified using methods described in (SI-1). The sample solutions were stored at 4 °C or −30 °C until further use. The concentration was calculated from the FAD and FMN molar absorption coefficient of ε_473_ = 9200 M^- 1^·cm^-1^ before use. The sample buffer was PBS (phosphate buffered saline) (pH 7.5), except for AppA_1-398_ and BlrP1, which were very unstable in PBS. The buffers of AppA_1-398_ and BlrP1 were 20 mM CHES buffer (pH 9.0, NaCl 150 mM, glycerol 10 % [v/v]), and 25 mM Tris-HCl buffer (pH 8.0, 300 mM NaCl, 5 mM MgCl_2_, 2 mM EDTA, 2 mM DTT and Glycerol 5% (v/v)), respectively.

### Absorption spectrum measurement

For absorption measurements, a diode array spectrophotometer (Agilent Cary 8454, Agilent Tech. Japan) was used. A quartz cell with an optical path length of 1 cm was used for the measurements, and the sample concentration was approximately 50 µM. To measure the absorption spectra in the light state, the sample solutions were irradiated with a blue light LED (center wavelength: 480 nm) (CHR-3S, NISSIN ELECTRONIC CO. Japan) from the top of the sample cell and confirmed that the spectra remained unchanged with prolonged exposure time (light steady state).

### Transient absorption measurement

In the transient absorption spectra measurements, a pulsed dye laser with a central wavelength of 458 nm (HyperDye 300, Luminomics, USA), pumped by an XeCl excimer laser (Lamda Physik, XeCl operation, Germany), was used for the excitation. A Xe lamp (Max-302, Asahi Spectra, Japan) was used as the probe light, and a CCD camera with a spectroscope was used for detection. The gate width of the CCD camera was set to 100 µs, and the delay time between the excimer laser trigger and the CCD camera trigger was varied using a delay generator to obtain the spectra at each time point.

To accurately measure the rate constants, the absorption changes were monitored at a single wavelength. The pulsed laser for the excitation was the above dye laser system, and the probe light was a blue LED light with a center wavelength of 480 nm. To monitor the change in wavelength around 490 nm, where many BLUF proteins show a large absorption change, a long-pass filter was placed between the LED light path and the sample solution (central wavelength:492 nm and halfwidth: 20 nm). Since the absorption spectra of the BLUF proteins BlrP1, YcgF, and PapB are red-shifted considerably, even in the dark state, different filters were used to cut off the wavelength below 498 nm (center wavelength: 498 nm and halfwidth: 25 nm). The spectrum of the probe light covers the entire red-shifted region of the BLUF absorption (Fig. S1).

The probe light transmitted through the sample solution was detected by a photomultiplier tube (Hamamatsu, R1477, Japan), and the time evolution of the transmitted light was recorded using an oscilloscope. The repetition rate of the pump pulse was between 0.2 and 0.005 Hz, depending on the dark recovery rate. For samples with extremely slow dark recovery rates (e.g., AppA), we measured them while stirring the sample to avoid multi-excitation of the protein. We obtained the average of several signals to improve the signal-to-noise ratio. A temperature controller was used for the temperature-dependent measurements. Unless otherwise noted, all measurements were performed at 25 °C.

## Results

### Slow absorption changes of BLUF proteins in milliseconds

The time profile of the absorption change in the millisecond time region, monitored at 492 nm upon photoexcitation of OaPAC, which is a multi-domain BLUF protein, is shown in Fig. 3(a). The initial rapid rise represents fast reaction dynamics, which could be similar to that reported previously.^19,24^ Subsequently, a slow rise component is clearly observed in the millisecond time region. The profile in this time range (I_TrA_(t)) is well reproduced by the sum of two exponential functions:

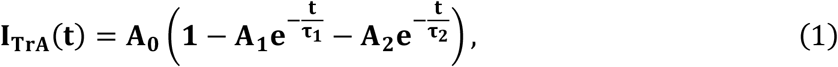

where A_0_ is the total amplitude, A_i_ (i = 1, 2) and τ_i_ (i = 1, 2; τ_1_ < τ_2_) are the amplitudes and time constants, respectively. The time constants and relative amplitudes are τ_1_ = 2.3 ms, τ_2_ = 36 ms, A_1_ = 0.023, and A_2_ = 0.33. To the best of our knowledge, this is the first clear observation of the absorption change in the millisecond time region for any BLUF protein.

**Fig. 3.**
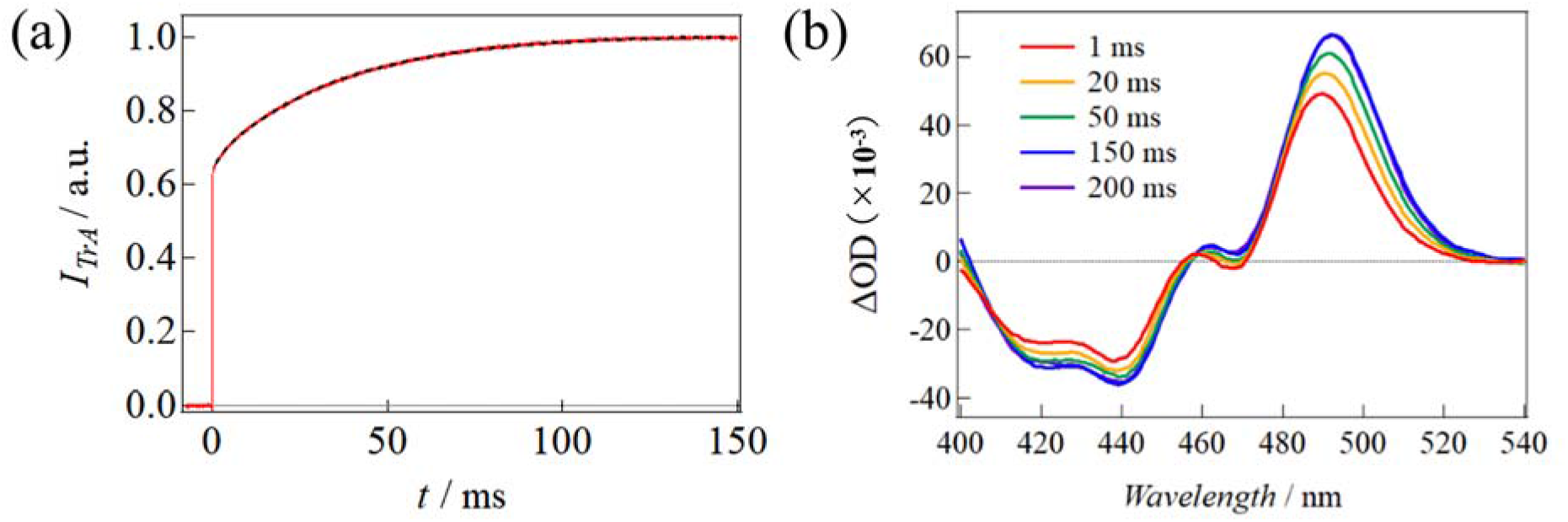
**(a)** Transient absorption signal of OaPAC in a millisecond time scale probed at 492 nm. The signal is normalized by A_0_ = 1. The fitting curves by eq. (1) are shown (dotted lines). **(b)** The transient absorption spectra in the millisecond time region of OaPAC.

Fig. 3(b) shows the transient absorption spectra of OaPAC upon photoexcitation. Within 1 ms, the absorption at 490 nm increased, reflecting the creation of red-shifted species. A further red-shift in the spectrum was observed until 150 ms. After 150 ms, no further absorption spectral change was observed. Hence, the observed slow rise of the TrA signal is due to the red-shift of the absorption spectrum, not due to enhancement of the absorbance with a constant red-shift of the product.

Since there has been no clear report on slow changes in the absorption spectrum for BLUF proteins (except the AppA_1-156_ construct described in the introduction), despite the many studies so far, the observation of the slow rising component was unexpected. Hence, to reveal the generality of this slow change in the BLUF proteins, the time profiles of the TrA signals for the multi-domain BLUF proteins (AppA_1-398_, OaPAC, BlrP1, YcgF), and the short BLUF proteins (SyPixD, TePixD, PapB), were measured and are shown in Fig. 4. For all proteins, the TrA signals exhibit rapid and slow rising phases on the millisecond time scale, although the relative amplitude of this slow phase varies depending on the specific protein. The time profiles of the signals are reproduced well by one or two exponential functions, as shown in Eq.(1) and Fig. 4. The parameters determined from the signal fittings are summarized in Table 1. These observations suggest that the slow absorption changes are common features of BLUF proteins.

**Table 1.**
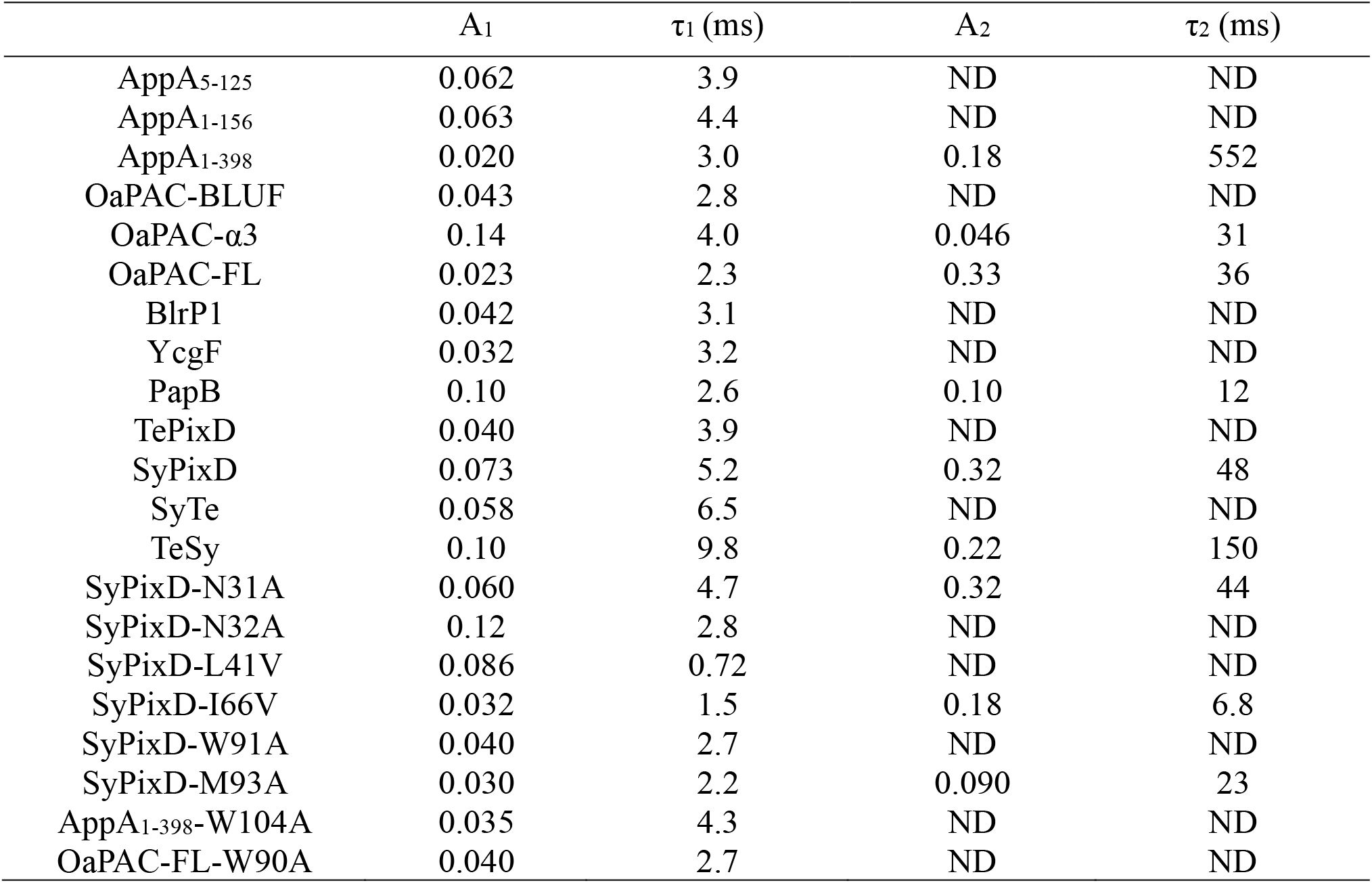
Time constants and their amplitudes of absorption changes in milliseconds. ND implies not detected.

**Fig. 4.**
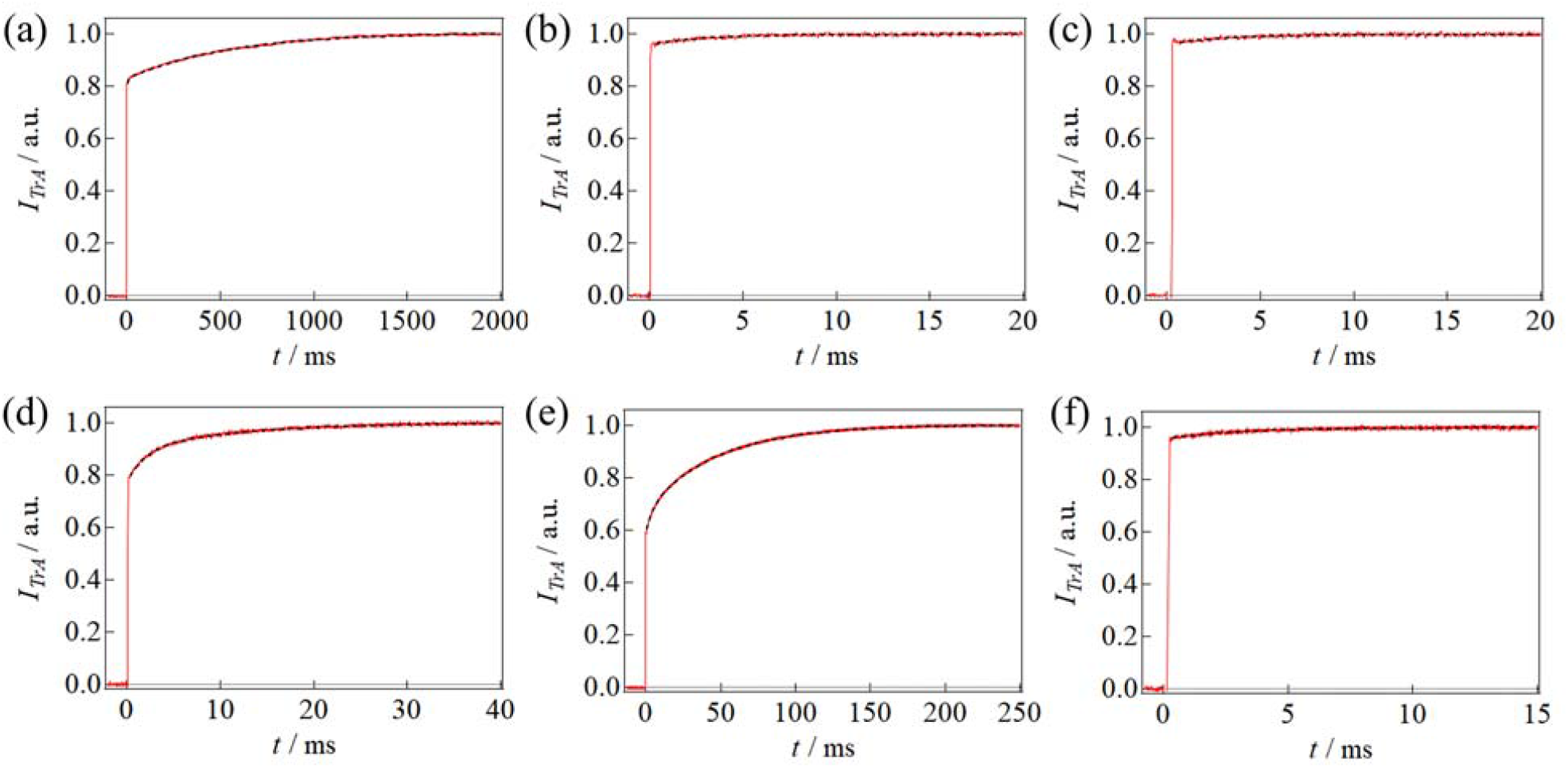
Transient absorption signal (red curves) in a millisecond time scale for (a) AppA_1-398_, (b) BlrP1, (c) YcgF, (d) PapB, (e) SyPixD, (f) TePixD. BlrP1, YcgF and PapB are probed at 498 nm and the others are probed at 492 nm: The fitting curves obtained by eq. (1) are shown (broken lines).

It should be noted that all BLUF proteins examined here exhibit multi-phase, ultrafast reactions, and a slow phase with a time constant of several milliseconds (τ_1_-phase). Some of them exhibit another additional phase with a time constant of tens of milliseconds (τ_2_-phase). For example, SyPixD exhibits two-phase dynamics in ms time range with τ_1_ = 5.2 ms and τ_2_ = 48 ms, while the ortholog protein TePixD exhibits one-phase dynamics with τ_1_ = 3.9 ms and A_1_ = 0.040. By examining the temperature dependence of the TrA of SyPixD, we concluded that these two reaction phases are sequential processes of the BLUF protein reaction (see SI-4 for details). This assignment is consistent with the TrA study of some mutants, as described below.

### Revisit of millisecond absorption change of AppA

The slow phase reactions of the BLUF proteins seemed to contradict the previous report on AppA_5-125_, which did not exhibit a millisecond absorption change.^19^ Thus, we re-examined the millisecond absorption changes of AppA_5-125_ and AppA_1-156_. The absorption spectra in the dark and light states of these constructs are almost identical (Fig. S2 (a)–(c)). Fig. 5 shows the TrA signals of these constructs. For AppA_1-156_, a weak change with τ_1_ = 4.4 ms was observed, which is consistent with the previous report.^24^ A similar slow-rising component was also observed for AppA_5-125_. AppA_5-125_ is a construct containing almost only the BLUF domain, and AppA_1-156_ contains the α3- and α4-helices in the C-terminal of the BLUF domain. Although the length of AppA_1-156_ is slightly longer, the difference does not to affect the slow dynamics.

**Fig. 5.**
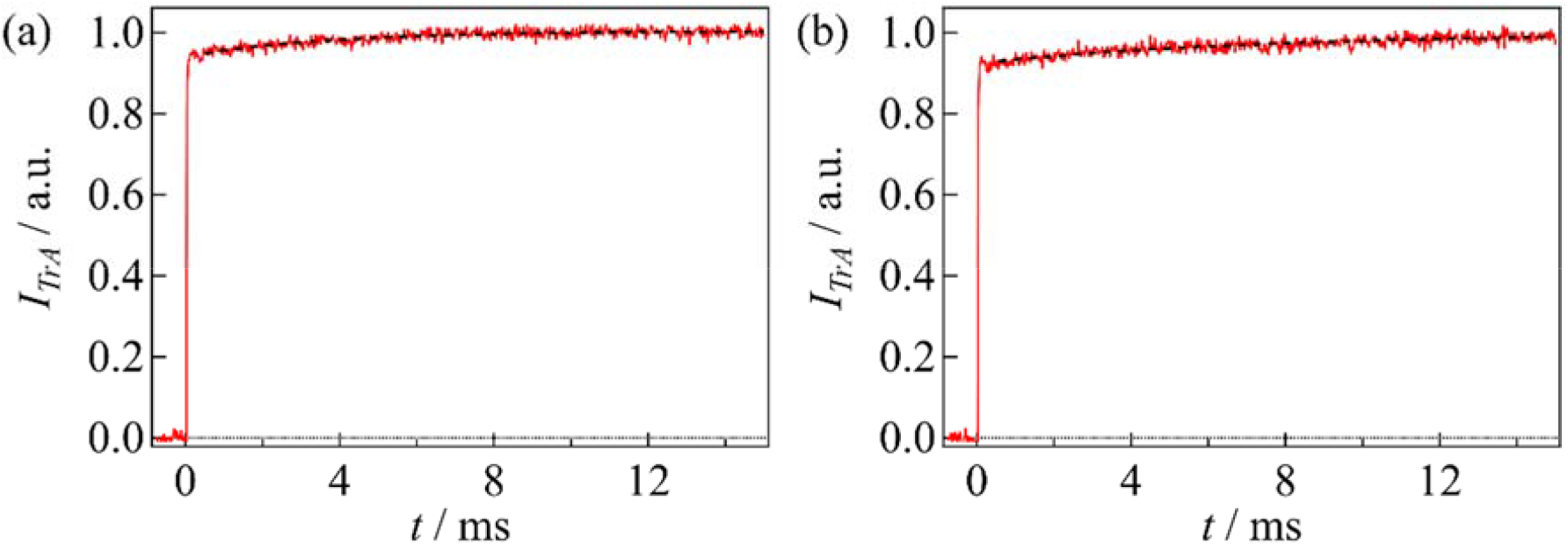
Transient absorption signals (red curves) of (a) AppA_5-125_ and (b) AppA_1-156_. The best fitted curves are shown in the black broken line.

### Effect of C-terminal domain

While the short constructs of AppA have only the τ_1_-phase, it should be noted that the profile of AppA_1-398_, which contains more bulky groups such as the SCHIC domain, shows a two-phase change (τ_1_ = 3.0 ms, τ_2_ = 552 ms) with relatively large amplitudes (Fig. 4(a) and Table 1). This result suggests that the slow change in the TrA signal depends on the size of the amino acid residues on the C-terminal side. To further study the importance of the C-terminal residues for the amplitude of the slow phase, the TrA signals of the C-terminal domain-deleted mutants of OaPAC were measured. OaPAC is a multi-domain BLUF protein, with a large C-terminus, which contains the α3-helix, and an AC domain (Fig. 6(a)).^32^ The time profiles of the TrA signals were monitored for a short construct OaPAC-BLUF (aa 1-96), which contains only the BLUF domain, and a construct OaPAC-α3 (aa 1-125), which contains the BLUF and the α3-helix of OaPAC, as well as the full-length OaPAC. The absorption spectra in the dark and light states of these constructs, and the full-length OaPAC, are almost identical (Fig. S1). This result indicates that the amount of the red-shift is determined only by the BLUF domain, and the C-terminus region does not contribute to the redshift. The TrA profile of OaPAC-α3 is reproduced by a bi-exponential function, eq. (1) with the time constants of 4.0 ms (A_1_ = 0.14) and 31 ms (A_2_ = 0.046). Although the bi-exponential behavior is the same as that for the full-length OaPAC, the relative amplitudes A_1_ and A_2_ are increased and reduced, respectively. The amplitude of the slower dynamics (A_2_) is further reduced for OaPAC-BLUF, and the signal is reproduced by a single-exponential function (A_2_ = 0 in eq.(1)) with a time constant of τ_1_ = 2.8 ms. These results suggest that the τ_1_-phase originates from the reaction of the BLUF domain, and that the slower τ_2_-phase represents the conformational change in the C-terminus region.

**Fig. 6.**
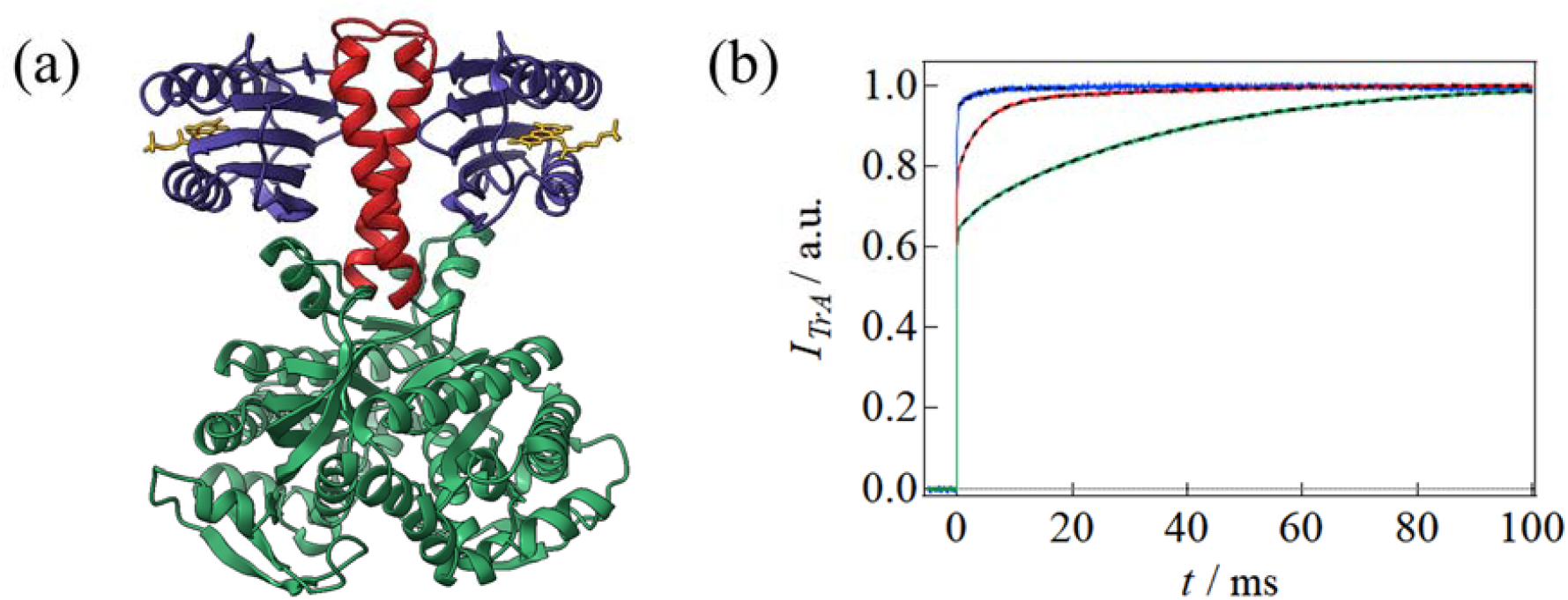
**(a)** Crystal structure of full-length OaPAC dimer. The 1-96 residues are colored by blue, 97-125 residues by red, and 126-366 residues by green. (b) The TrA signals of short OaPAC constructs; OaPAC-BLUF (blue), OaPAC-α3 (red), and full-length OaPAC (green). The best fitting curves by eq. are shown by the broken lines.

### Effect of C-terminal residues on slow dynamics of TePixD and SyPixD

It is interesting to note that the amplitudes of the slow-rise components are very different between SyPixD and TePixD, although SyPixD and TePixD are orthologous proteins and have very similar decameric structures (Fig. 7(a)). The TrA profile of SyPixD is well reproduced by the two time constants (τ_1_, τ_2_) with a large amplitude, whereas that of TePixD is reproduced by a τ_1_-phase with a small A_1_. The primary sequences of SyPixD (1-150) and TePixD (1-143) show that the BLUF domains of these proteins have a relatively high identity (47/98; 48%), but the C-terminal regions are significantly less homologous (14/46; 30%). Hence, we speculate that the differences in these slow phases result from the difference in the C-terminus. To examine this possibility, we prepared two chimeric mutants, a SyTe mutant, which consists of the BLUF domain (1-102) of SyPixD with the C-terminal region of TePixD (103-143), and a TeSy mutant, which consists of the BLUF domain (1-102) of TePixD with the C-terminal region of SyPixD (103-150). The TrA signals were measured and are shown in Fig. 7(b). Interestingly, the τ_2_-phase in the TrA signal of TeSy is significantly stronger than that of TePixD, and the profile exhibits the two-phase dynamics, whereas SyTe exhibits a weak rise component with only the τ_1_-phase (Fig. 7(b)). These results indicate that the different profiles for SyPixD and TePixD are caused by the different lengths and/or characters of the C-terminal regions, although the absorption spectrum in the visible wavelength region is determined by the residues close to the chromophore flavin.

**Fig. 7.**
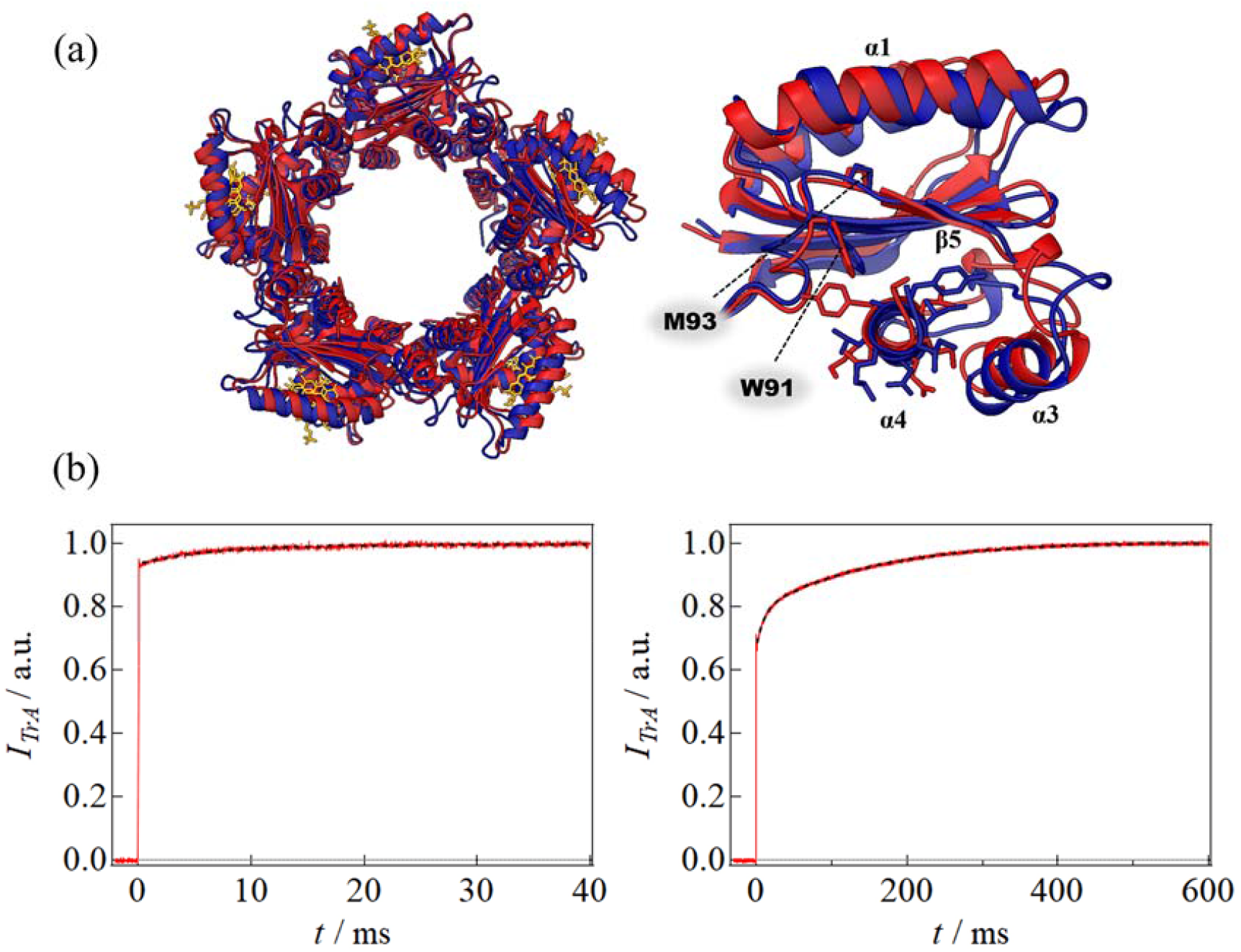
**(a)** Overlapped crystal structures of SyPixD (blue, 2HFN) and TePixD (red, 1X0P). Decamer (left), monomer (right). **(b)** TrA signals of SyTe (left) and TeSy (right).

### Contributions of residues in the BLUF domain of SyPixD

Since the absorption in the visible wavelength is determined by the residues close to the flavin, the slow absorption change must be due to a slow conformational change around the flavin. We next investigated the residues that may cause the slow change using point mutations around the chromophore of SyPixD. We chose SyPixD because the amplitude of the millisecond absorption change is large and the roles of some residues in the BLUF domain have been studied previously.^29,31,18^ The positions of residues for the point mutation in this study are shown in Fig. 8(a).

**Fig. 8.**
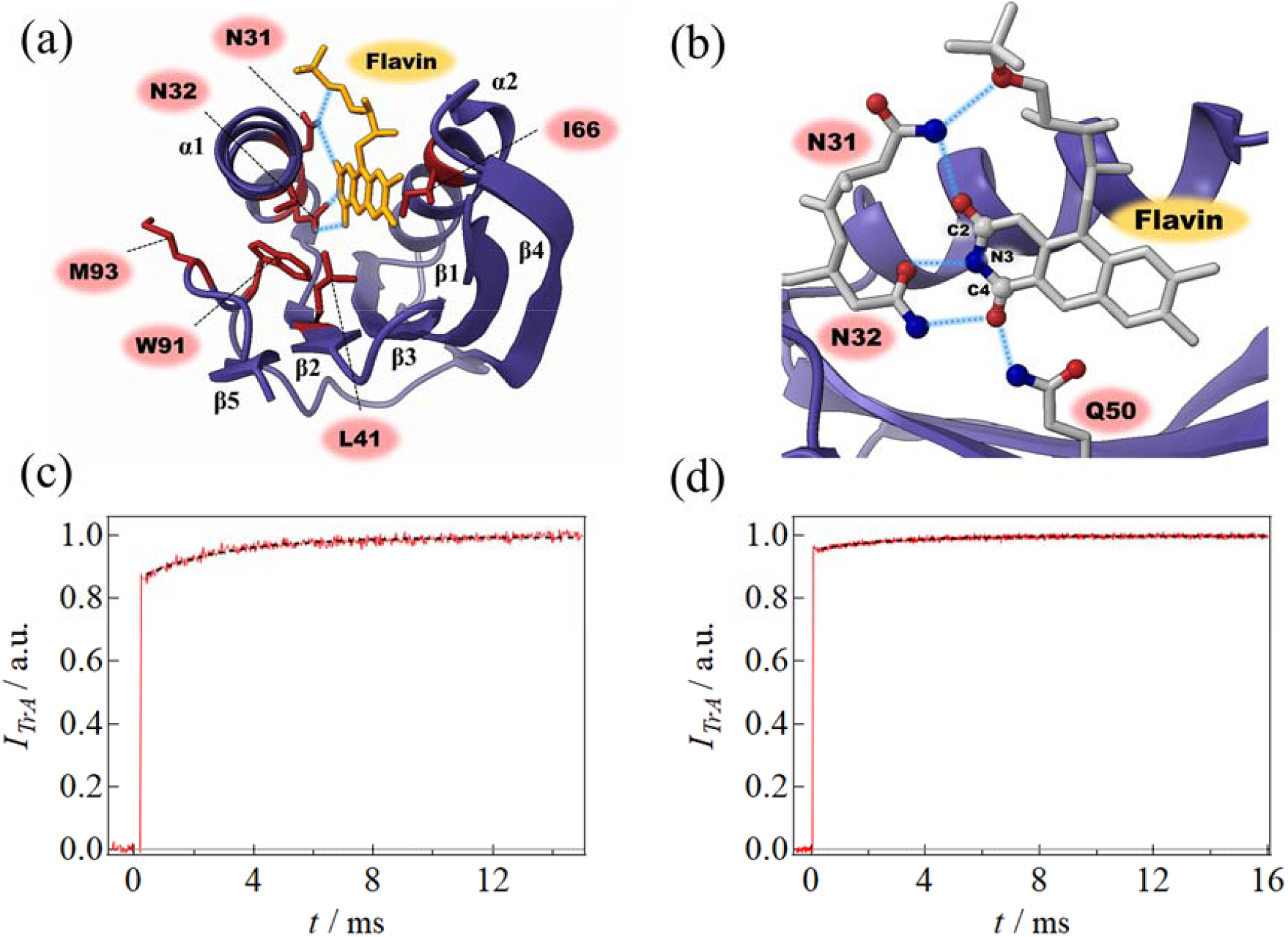
(a) The positions of amino acid residues of SyPixD used for mutation studies. (b) Hydrogen bond network around chromophore. Oxygen atoms are in red and nitrogen atoms are in blue. (c) TrA signal of SyPixD-N32A and (d) SyPixD-W91A.

#### (a) SyPixD-N31A and N32A

Both N31 and N32 of SyPixD are located on the α1-helix of the BLUF domain. N32 is conserved in all the BLUF proteins (Fig. 2(b)), but N31 is not. The side chains of N31 and N32 are hydrogen-bonded to C_2_=O of the flavin, and to N3-H and C_4_=O, respectively (Fig. 8(b)).^4^ One of the causes of the spectral red-shift upon photoexcitation is considered to be the formation of the hydrogen bond between these residues and C_4_=O.^16,17^ The absorption spectrum of SyPixD-N32A shows a red-shift of 4 nm (from 440 nm to 444 nm), which is significantly smaller than that of the wild type (WT) SyPixD (14 nm). A similar reduction in red-shift was also reported for TePixD-N32A.^6^ The TrA signal of SyPixD-N32A exhibits a weak τ_1_-phase, with a time constant of τ_1_ = 2.8 ms (A_1_ = 0.12), but the τ_2_-phase is absent. Hence, SyPixD-N32 is an important residue not only for producing the red-shift, but also for its slow dynamics. In particular, this residue is one of the origins of the τ_2_-phase, which is caused by the conformational change in the C-terminal group. On the other hand, the slow rise of SyPixD-N31A is almost the same as that of WT (SI-5).

#### (b) SyPixD-L41V and I66V

L41 and I66 of SyPixD are located on the β2-strand and α2-helix of the BLUF domain, respectively, and are highly conserved among the BLUF proteins (Fig. 2(b)). The alkyl groups in the side chains are located in the vicinity of the isoaloxazine ring of the flavin and are considered to be responsible for retaining the flavin in the BLUF domain through the hydrophobic interactions. Hence, by replacing the bulky alkyl groups of L41 and I66 with smaller ones, extra space is created around the flavin to expand in the movable space. SyPixD-L41V exhibits two-phase slow dynamics τ_1_ = 0.72 ms and τ_2_ = 11 ms (SI-5, Fig. S4 (b)). Similarly, SyPixD-I66V shows two-phase slow dynamics with τ_1_ = 1.5 ms and τ_2_ = 6.8 ms (SI-5, Fig. S4 (c)). Both these rates are several times faster than that of WT (5.2 ms (τ_1_) and 48 ms (τ_2_)).

#### (c) SyPixD-W91A and M93A

W91 and M93 of SyPixD are located in the loop region close to the β5-strand of the BLUF domain and are conserved in many BLUF proteins (Fig. 2(b)). W91 is one of the most interesting residues because it takes two different conformations synchronized with M93 in the crystal structures.^4^ These residues have been proposed to be important for the intramolecular signal transduction, although their importance is controversial, as described in the introduction.^14^ The transient absorption of SyPixD-W91A showed a one-phase change, with τ_1_ = 2.7 ms and A_1_ = 0.04. On the other hand, SyPixD-M93A retained the same two-phase changes as the WT, although the amplitudes are decreased (SI-5, Fig. S4 (d)).

## Discussion

### Origin of τ_1_-phase

In this study, we found that all BLUF proteins examined here exhibit slow (milliseconds) changes in the absorption in the visible wavelength region. The slow components are expressed well by single- or bi-exponential functions (eq.(1)). Importantly, the τ_1_-phase is well conserved in all the BLUF proteins. Compared to A_2_, A_1_ is not very sensitive to proteins and mutations. Furthermore, the time constants τ_1_ are similar among BLUF proteins. Therefore, it is conceivable that the τ_1_-phase is a common reaction in the BLUF domain and may be a key step in the subsequent conformational change of the protein.

The origins of the spectral red-shift upon photoexcitation of the BLUF proteins have been studied extensively.^16,17,18,33^ The red-shift is caused by the changes in the hydrogen bonds around the flavin. For example, N31 of SyPixD interacts with the C2=O bond of the flavin via a hydrogen bond (Fig. 8(b)).^4^ Hence, it is possible that the slow changes in the position of N31 could cause the τ_1_-phase in the absorption spectrum. However, the rates and amplitudes of the slow components of SyPixD-N31A are similar to those of WT SyPixD. Therefore, we suggest that the hydrogen bond to C2=O does not contribute to the slow change.

It has been suggested that N32 and Q50, which are hydrogen bonded with the C_4_=O of the flavin (Fig. 8(b)), are responsible for the red-shift,^6^ because SyPixD-Q50A does not form a red-shift state upon light illumination, and the red-shift of SyPixD-N32A is significantly reduced. Indeed, X-ray crystallographic structures of OaPAC in the dark and light states showed that the hydrogen bond distances between C_4_=O and the N32 and Q50 side chains of flavin decrease by 0.3 Å and 0.5 Å, respectively, in the light state.^9^ The decrease in the hydrogen bond distance reduces the double-bonded nature of C_4_=O and causes the delocalization of the π-electrons in the flavin, which leads to a red-shift in the absorption spectrum. It was found that the amplitude of the τ_2_-phase, A_2_ is significantly decreased for SyPixD-N32A compared with WT SyPixD. Hence, N32 is certainly important for the τ_2_-phase. According to the SyPixD computational study,^30^ changes in the hydrogen bond between Q50 and the C4 carbonyl of flavin affects the β5-strand via N32, and our results support this idea. On the other hand, the amplitude of the τ_1_-phase, A_1_, does not change significantly with the N32A mutation. Since N32 and Q50 mainly cause the spectral red-shift, we consider the τ_1_-phase to occur via the movement of Q50 on the β3 strand.

The rates of absorption changes (τ_1_ and τ_2_) are faster in the SyPixD-L41V and I66V mutants (SI-5). This result may be due to the weakening of the hydrophobic interaction with the isoalloxazine ring of the flavin. The void space around the flavin becomes larger by replacing the bulky side chains with smaller ones. These results suggest that the absorption change in the millisecond time scale may be caused by a slight shift of the chromophore flavin toward the N32 side chain. Therefore, we suggest that the τ_1_-phase is strongly correlated with the Q50 motion on the β3 strand and that the flavin motion in the pocket is responsible for the spectral change.

### Origin of τ_2_-phase

We found that the slower dynamics (τ_2_-phase) are sensitive to the C-terminal region, at least for AppA, OaPAC, and Sy(Te)PixD. This point is important, because of the significance of the C-terminal region of the BLUF proteins for the biological functions has frequently been pointed out.^7,9,34,35^ For example, the crystal structure analysis of OaPAC shows that photoreaction induces a conformational change in the C-terminal adenylate cyclase domain via the α3-helix.^9^ Furthermore, studies using a chimeric mutant of SyPixD, in which the BLUF domain is connected to the C-terminal α3- and α4-helices of PapB, showed signal transformation from the BLUF domain through the C-terminal helix.^35^ In this respect, it is important to note that the τ_2_-phase is an indicator of the conformational change of the C-terminal domain, which is far from the chromophore. From these observations, one may speculate that the conformational changes in the C-terminus region somehow influence the conformation of the residues in the BLUF domain. However, we believe this is not the case, as explained below.

In Fig. 9, the amplitudes of the slow dynamics, A_1_, A_2_, and A_1_ + A_2_, are plotted against the amount of the red-shift of the absorption spectrum, *E*_red_, which is defined by the energy difference between the half absorbances of the peaks of the absorption spectra in the light and dark states (SI-3). This plot and Fig. S1 show that the steady-state absorption spectra of AppA and OaPAC, with or without the C-terminal groups are almost identical both in the dark and light states. In other words, the C-terminal groups do not influence the hydrogen bond network around the flavin, and the absorption spectrum is determined only by the BLUF domain. Since the BLUF domains of AppA (AppA_5-125_) and OaPAC-BLUF do not have the τ_2_-phase, the red-shifts of these proteins without the C-terminal group should complete within the τ_1_-step. By connecting the C-terminal domain to the BLUF domain of these proteins, the τ_2_-phase appears without changing *E*_red_. Hence, we should consider that a part of the ultrafast red-shift process of the BLUF domain, that is, the conformational change around the flavin, is stopped before the final state occurs, and that the final state is established slowly in the τ_2_-phase. During this τ_2_-step, the conformational change of the C-terminal group allows the relaxation of the conformation in the BLUF domain. Interestingly, among the wild type-BLUF proteins, a correlation between *E*_red_ and A_2_ is noticeable, that is, the larger the *E*_red_, the larger the A_2_. A larger *E*_red_ indicates that the changes in the hydrogen bonds around the flavin are larger for these proteins, which is mostly explained by the increase in A_2_. Hence, the observed correlation probably indicates that the ultrafast changes in the hydrogen bonding and τ_1_-phase are similar among the BLUF proteins, and the τ_2_-phase varies depending on the protein.

**Fig. 9.**
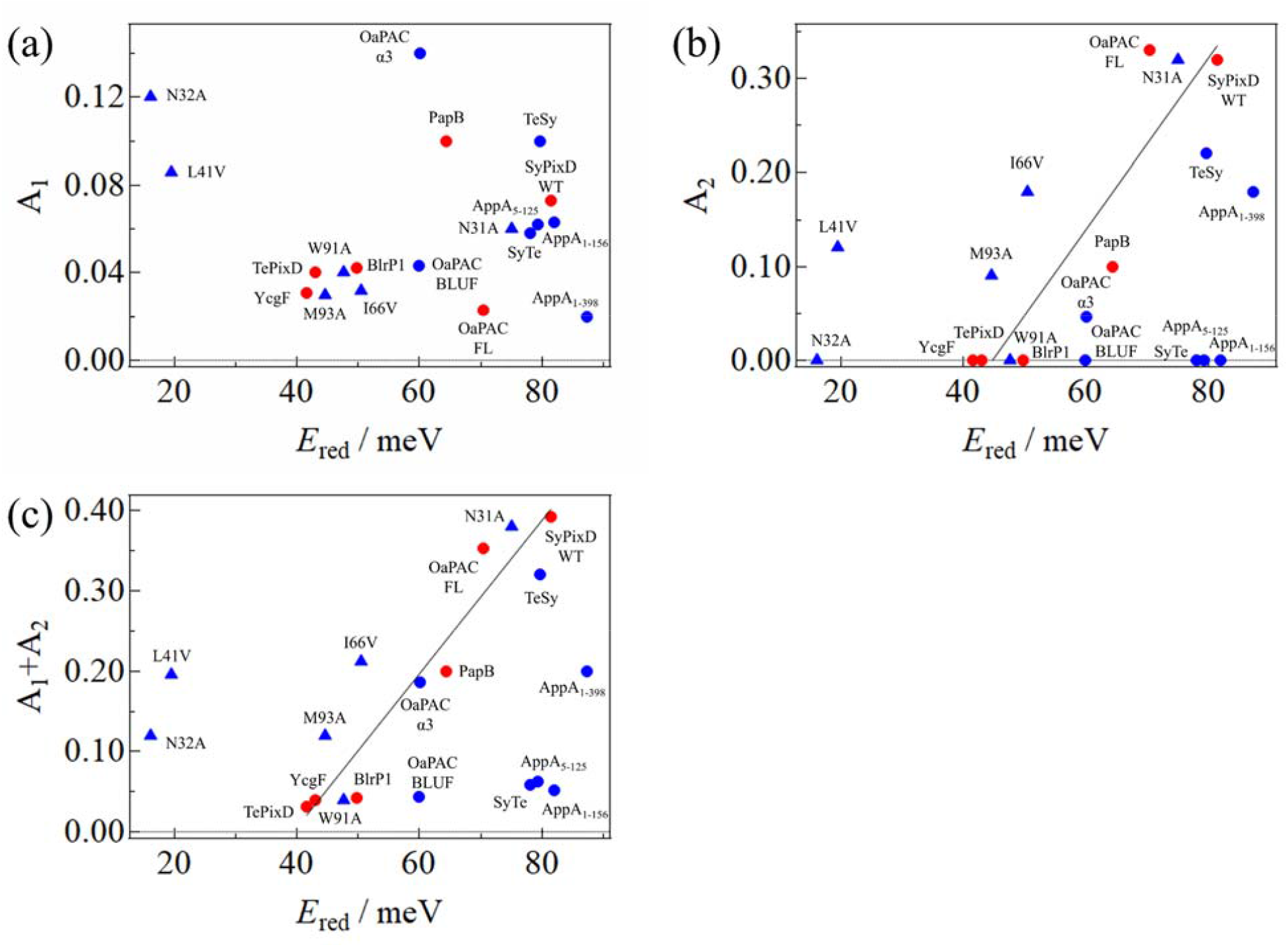
Plots of (a) A_1_ vs *E*_red_, (b) A_2_ vs *E*_red_, (c) A_1_ + A_2_ vs *E*_red_. WT proteins are shown in red, and the others in blue. SyPixD mutants are indicated by triangles. The black straight lines in (b) and (c) are the best fitted lines by a linear function for the WT proteins.

For SyPixD-N32A and W91A, the τ_2_-phase disappeared. Therefore, these residues are important for linking the C-terminal structural changes to the local flavin environment. In particular, W91 interacts directly with the C-terminal cap helix in the Trp_out_ conformation (Fig. 7(a)),^4^ and its importance in signal transduction has been widely studied. Fluorescence quenching measurements in SyPixD showed that W91 is partially exposed to the solvent side and its exposure increased slightly with light.^29^ This result suggests a conformational change of W91 in the light state. It has been speculated that W91 is responsible for propagating conformational changes from around the chromophore to the C-terminal region.^1^ Hence, we consider that the loss of the τ_2_-phase in SyPixD-W91A may be due to the lack of a connection. To further support the importance of the Trp residue, the slow reaction of W/A mutants of AppA_1-398_ and OaPAC were measured and shown in Fig. S1-6. The profiles did not exhibit the τ_2_-phase. These observations clearly indicate the importance of this residue. In fact, FTIR studies of AppA and its mutant, AppA-W104A (corresponding to SyPixD-W91A), revealed that the signals of the mode associated with C_4_=O of the flavin ring and the amide band of β-sheet disappear in AppA-W104A.^36^ Moreover, X-ray crystal structure analysis of OaPAC in the dark and light states shows that W90 is oriented outside the BLUF domain in the dark state, and that the orientation changes slightly with light.^9^ These observations are consistent with our findings.

W91 is not conserved in BlrP1 and YcgF. Interestingly, these proteins do not exhibit the τ_2_-phase, which supports the speculation described above, that is, the slow spectral change is related to the movement of the Trp residue. Previous transient grating (TG) measurements have revealed that the two proteins (BlrP1 and YcgF) also show millisecond conformational changes, despite the absence of Trp and the τ_2_-phase.^37,38^ Hence, in these cases, the slow conformational change may be caused by the τ_1_-phase through different signaling pathways. NMR measurements of BlrP1 indicate that the blue light induces conformational changes in α3-and α4-helices extending from the C-terminus of the BLUF domain.^39^ In the case of BlrP1, the C-terminal cap helix is docked on the β-strands of the BLUF domain, like in other BLUF proteins, such as PixDs,^4,6^ AppA,^34^ OaPAC,^9^ bPAC,^40^ BlrB,^5^ and BlsA,^41^ but the orientation of the helix relative to the β-strands is different in BlrP1 from the others (parallel for BlrP1 and perpendicular for the others).^42^ As a result, the distance between the position of Trp, which is replaced by Thr in BlrP1, and the cap helix is longer in BlrP1 than the other proteins. These structural differences suggest that BlrP1 has a different signal transduction mechanism.

TePixD does not exhibit the τ_2_-phase, despite having Trp. Interestingly, however, the transient absorption signals showed that TeSy exhibits a two-phase change with a large intensity, whereas SyTe shows a one-phase change with a small intensity. The characteristic slow absorption change is completely opposite to that of the WT, indicating that the character of the C-terminus determines the appearance of the τ_2_-phase. The crystal structures of SyPixD and TePixD are very similar, but the residues around W91, and the C-terminal helix under the β5 strand are different (Fig. 7(a)). These differences likely affected the movement of the W91. Therefore, the presence of W91 is necessary but not sufficient, and the nature of the C-terminus determines the appearance of the τ_2_-phase.

The significant decrease in the relative amplitude of the τ_2_-phase (A_2_) for M93A of SyPixD suggests suppression of signaling to the C terminus by this mutation. M93 is located near Q50, such that these two residues form a hydrogen bond and are highly conserved in other BLUF proteins. Therefore, M93 is also thought to be involved in connecting the movement of Q50 to the conformational change of the β5 strand. Indeed, a mutation on M93 (M93A) inhibits the light-induced change in the amide-II region in the FTIR difference spectrum,^31^ indicating that M93 is an essential residue for transmitting light information to the β5 strand. The importance of M93 is further evident from the fact that *Synechocystis* strain expressing the M93A mutant exhibits a negative phototaxis, in contrast to the positive phototaxis of the WT strain.^31^ These observations are consistent with our above suggestion.

A time-resolved FTIR study on AppA_5-125_ has been reported,^36^ and it was shown that the secondary structure change occurs with a time constant of 1-5 μs, which is much faster than our observed rates. This study was performed on a short construct of AppA, which does not exhibit the τ_2_-phase. We may consider that the absence of C-terminal residues accelerates the conformational change of the β5 strand to a few microseconds, or that the conformational change is not completed within the observation time window of the time-resolved FTIR study (up to 1 ms). Indeed, the time-resolved IR difference spectrum recorded at 20 μs does not completely overlap with the steady-state IR difference spectrum. Thus, it is possible that further structural changes occur on the millisecond timescale, which have been observed in this study.

### Signaling pathway of BLUF proteins

Previously, the global conformational changes of many BLUF proteins have been detected by the TG technique.^37,38,43,44,45^ For example, the time constants of the conformation changes far from the chromophore for PapB, BlrP1, SyPixD, and TePixD have been reported to be 24 ms,^45^ 21 ms,^38^ 350 ms,^46^ and 4 ms,^44^, respectively. It is interesting to note that the time constant of PapB (24 ms) are relatively close to those observed from the slow absorption change (τ_2_ = 12 ms). Hence, we consider that the conformational change of PapB detected by the TG method induces conformational changes in the hydrogen bond network around the flavin. On the other hand, the rates of the conformational changes of BlrP1 and SyPixD are much slower than those from the absorption changes. In this case, these changes may be triggered by the τ_1_-phase and/or τ_2_-phase, and do not influence the hydrogen bond network around the flavin.

Based on the findings described above, we propose a signaling mechanism for BLUF proteins (Fig. 10). The ultrafast reaction occurs within nanoseconds due to the transfer of electrons and protons between flavin and its neighboring Y8, and the orientation change of Q50.^47^ Following this localized change, the flavin moves through the binding pocket (τ_1_-phase), of which the rate constants are nearly identical among BLUF proteins and dependent on the size of the space in the pocket. Subsequently, in the case of SyPixD, W91 and N32 move together to complete the red-shift (τ_2_-phase). The τ_2_-phase may accompany the conformational change of the β5 strand, leading to the following global reaction observed by the TG method (dissociation of the super complex in the case of SyPixD).^43^ It is important to note that the size and nature of the C-terminal residues of the BLUF domain strongly influence the τ_2_-phase. This indicates that the interaction with the C-terminal region slows down the movement of W91, and the τ_2_-phase clearly appears. Deletion of the C-terminus increases the flexibility of W91 and completes the reaction synchronously with the τ_1_-phase.

**Fig. 10.**
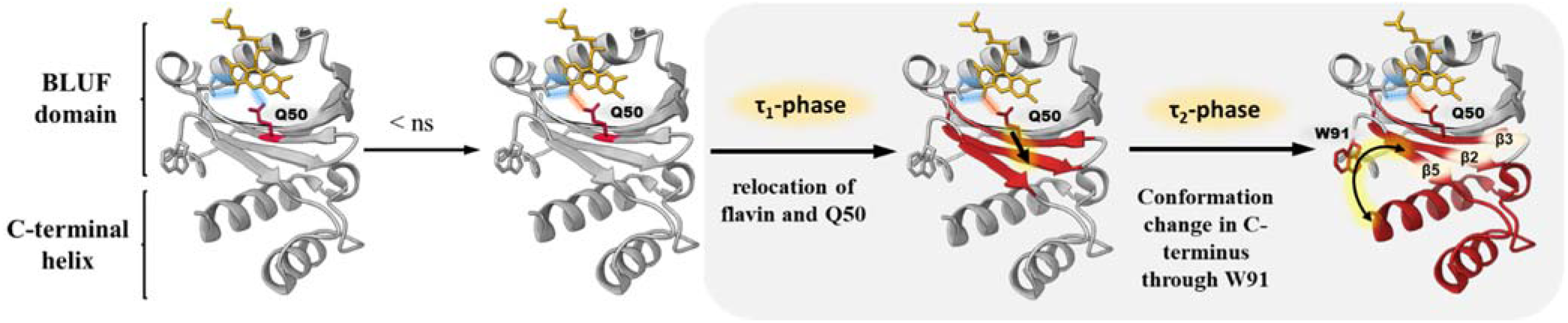
Reaction scheme of BLUF proteins, e.g., SyPixD. The blue line represents hydrogen bonds in the dark state. Within a nanosecond, the orientation of the side chain of Q50 changes, and a new hydrogen bond is formed with the C_4_=O of the flavin (shown by the red line). In the τ_1_-phase, the distance between Q50 and flavin slightly decreases, probably due to movement of β-strands. In the τ_2_-phase, W91 transmits the movement of β5 to the C-terminus, resulting in a further reduction of the distance between flavin and Q50 through the β-sheet. The structure in red indicates the area of the conformational change, and the black arrows indicate the signaling pathway.

### Summary

In this study, we found that the absorption in the visible wavelength range changes in the millisecond time range for all the BLUF proteins we investigated. Commonly, the slow dynamics are expressed well by a bi-exponential function with faster (τ_1_-phase) and slower (τ_2_-phase) processes. The origins of these slow changes were investigated. The amplitude of the τ_2_-phase is strongly dependent on the size and nature of the C-terminal groups, as demonstrated by the truncated mutants of OaPAC and chimeric mutants of SyPixD and TePixD. We also found that the conformational change of the C-terminal group affect the chromophore absorption spectrum through Asn32 and Trp91 of SyPixD. In particular, Trp on the β5 strand was found to be related to the τ_2_-phase in OaPAC and AppA. We suggest that a part of the ultrafast red-shift process of the BLUF domain, that is, the conformational change around the flavin, is stopped before the final state, and the final state is established slowly in the τ_2_-phase. During this τ_2_-step, the conformational change of the C-terminal group allows the relaxation of the conformation in the BLUF domain. Since the conformations of the C-terminal groups of the BLUF proteins usually control the biological activities, the observed slow dynamics should be an important indicator for understanding the whole picture of the photoreaction that regulates the function of BLUF proteins. In contrast, the τ_1_-phase is observed for all BLUF proteins, including the mutants. Hence, this phase is a common feature of the BLUF proteins.

## SUPPORTING INFORMATION

Expression and purification of sample proteins (SI-1), Absorption spectra and probe light profiles (SI-2), Definition of Ered (SI-3), Slow rise components of SyPixD at various temperatures (SI-4), A Transient absorption signals of SyPixD mutants (SI-5), Transient absorption signals of W/A mutants of AppA1-398 and OaPAC (SI-7).

## ACKNOWLEDGEMENTS

This work was supported by a Grant-in-aid for Scientific Research on Innovative Areas (research in a proposed research area) (Nos. JP20107003, and JP25102004) and a Grant-in-aid for Scientific Research (17H03008, 21H01885, 21K19218 to M.T. and 18H045522, 20H04708 to Y.N.) from MEXT/JSPS.

The authors have no conflicts to disclose.

## Notes

### Competing Interest Statement

The authors have declared no competing interest.

