## Supporting Information for "Slow conformational changes of blue light sensor BLUF proteins in milliseconds"

### SI-1. Expression and purification of sample proteins

For expression of the BLUF proteins, pET28 was used as the vector (pUG57 for BlrP1), and BL21(DE3) pLysS (BL21(DE3) for AppA and PapB, and Rosetta2(DE3) for BlrP1) was used for the *E. coli* strain. AppA is fused with C-terminal his-tag, and the others are with N-terminal his-tag. The his-tagged proteins were prepared by the following procedure except for AppA and BlrP1. *E. coli* carrying the plasmid encoding the gene of the target protein was cultured in LB medium at 37 °C until OD600 reached 0.5~0.6. IPTG was added to the medium to a final concentration of 0.1 mM and incubated at 18 °C for about 20 hours. Cells were harvested by centrifugation at 4000 g for 15 min at 4 °C and suspended in PBS buffer with DnaseI and excess FAD. After disruption of the cells by sonication, the homogenate was centrifuged at 20000 g for 1 h at 4 °C. The His-tagged protein was purified from the supernatant by Ni-affinity column chromatography (HisTrap HP, GE Healthcare). After the medium was exchanged to PBS buffer (pH7.5) using a desalting column, the N-terminal His-tag was cleaved off by turbo3C protease for SyPixD. The cleaved His-tag polypeptide and the uncleaved fusion protein were trapped by passing over the Ni column. The flow-through from the column was pooled and concentrated as a final purified sample solution. The appropriate cleavage and purity of > 95% were confirmed by SDS-PAGE. For AppA and BlrP1, the previously reported method was used.[1,2]

### SI-2 Absorption spectra and probe light profiles

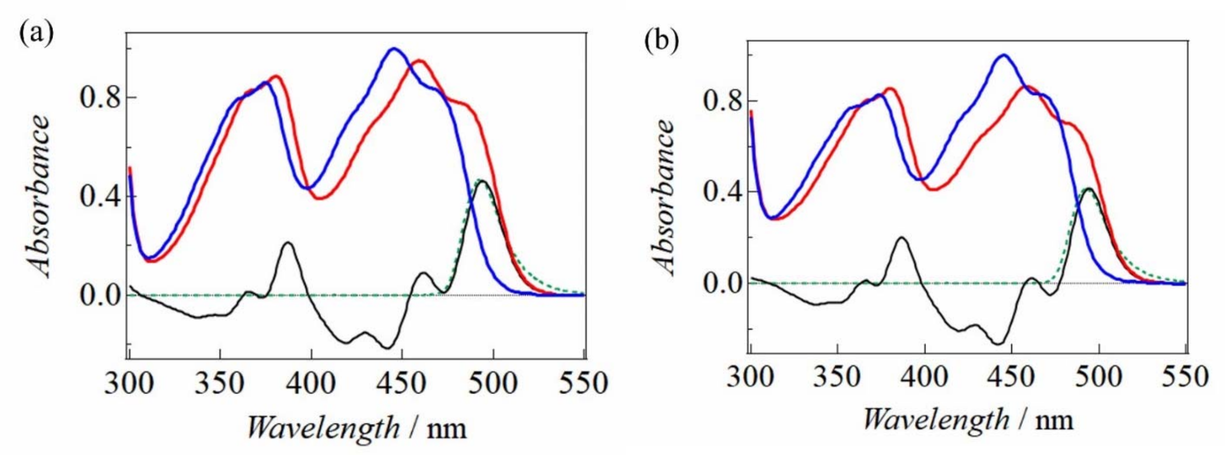

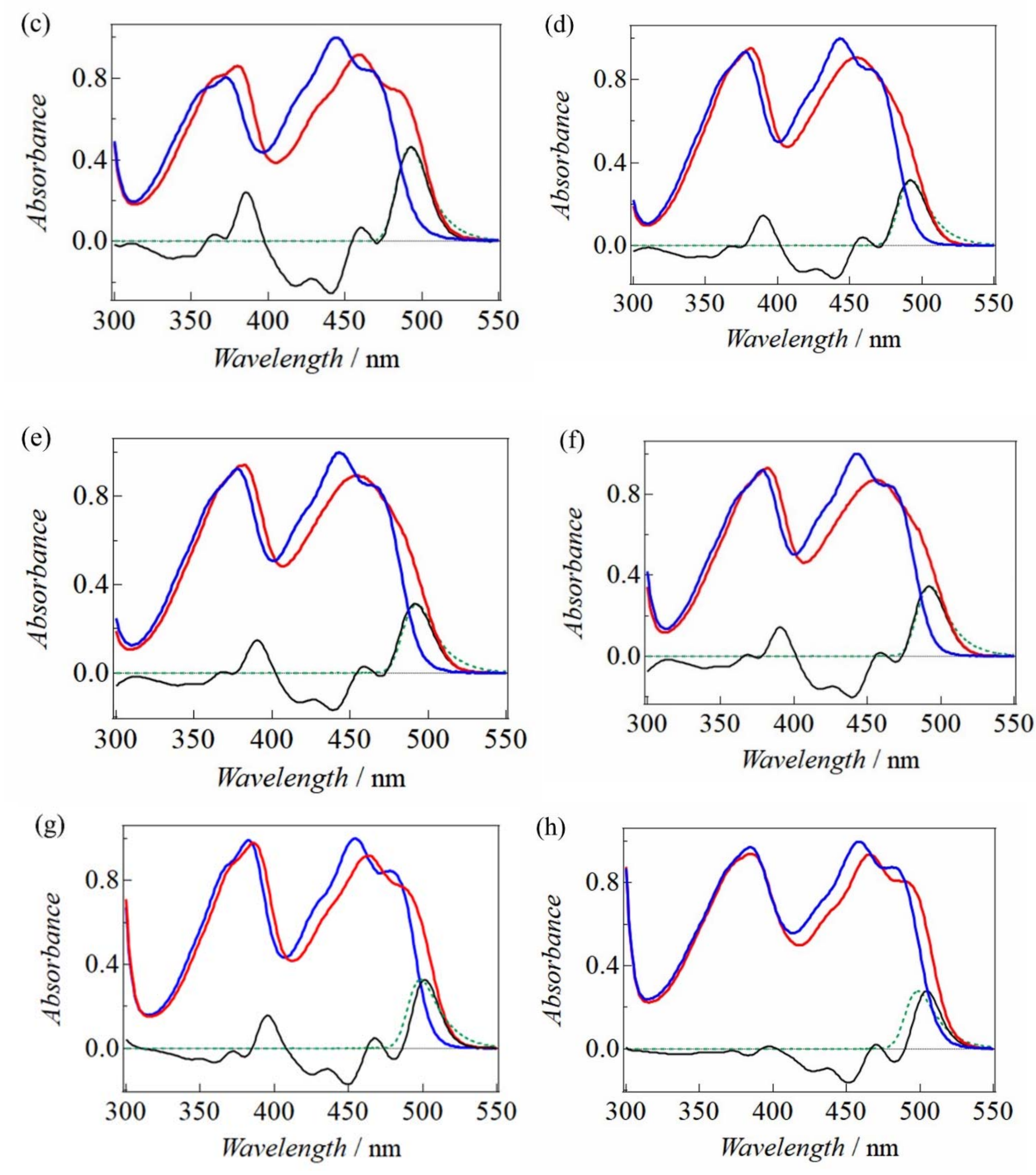
(c)
(d)
(e)
(f)
(g)
(h)
Absorbance
0.0
0.4
0.8
300
350
400
450
500
550
Wavelength / nm

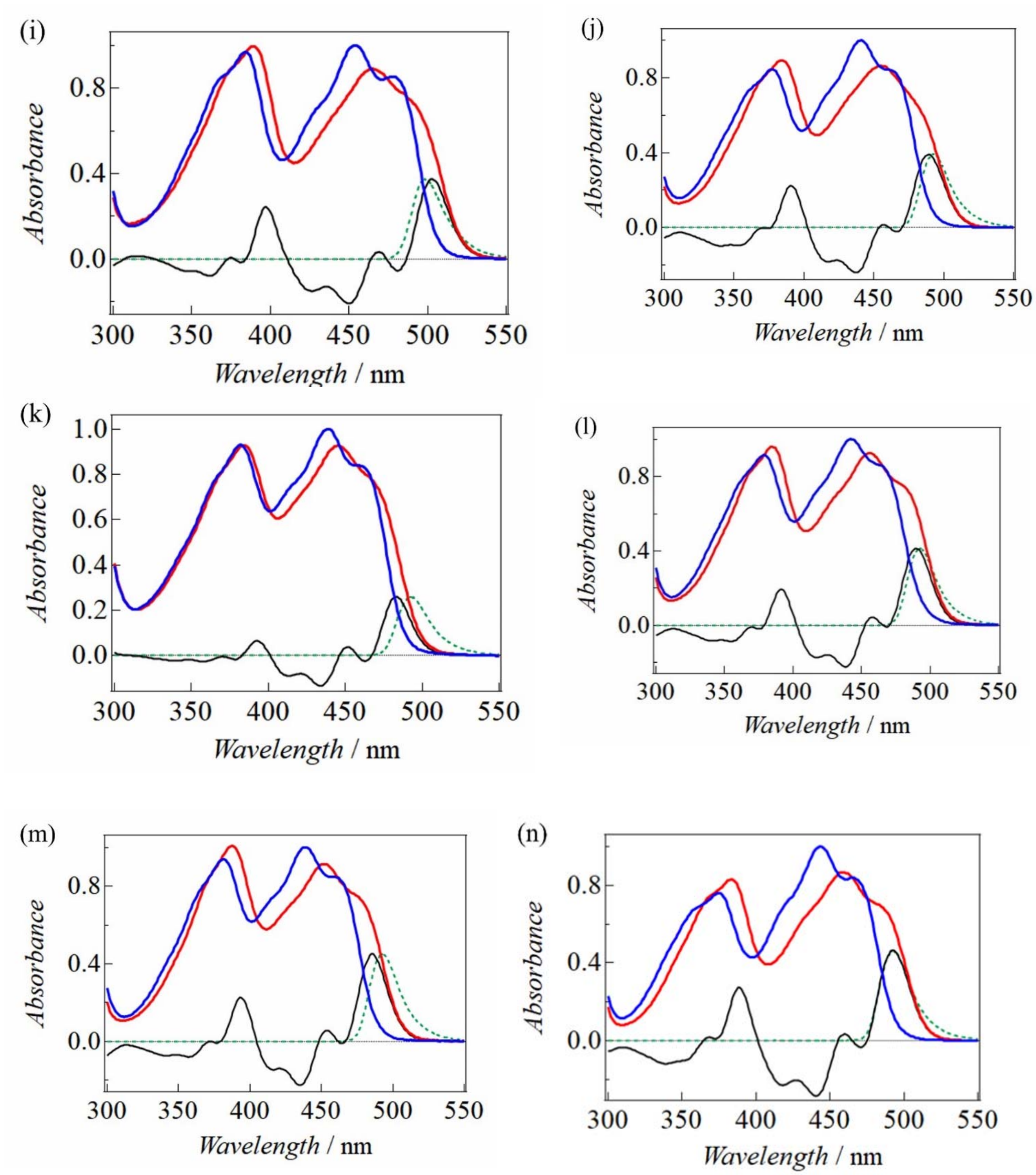
(i)
(j)
(k)
(l)
(m)
(n)
Absorbance
Wavelength / nm
300
350
400
450
500
550
0.0
0.2
0.4
0.6
0.8
1.0

(o)

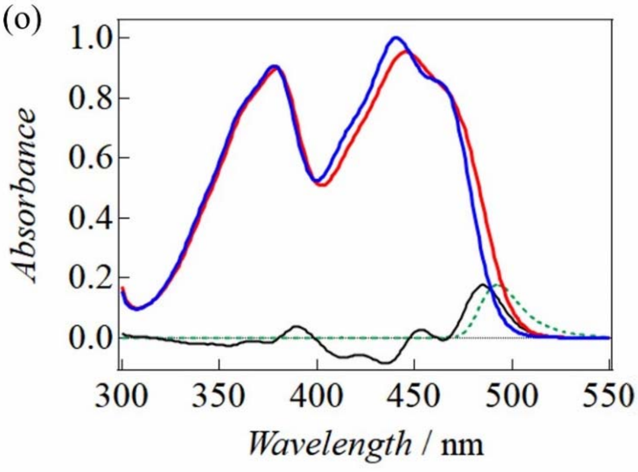
1.0
0.8
0.6
0.4
0.2
0.0
Absorbance
300
350
400
450
500
550
Wavelength / nm

(p)

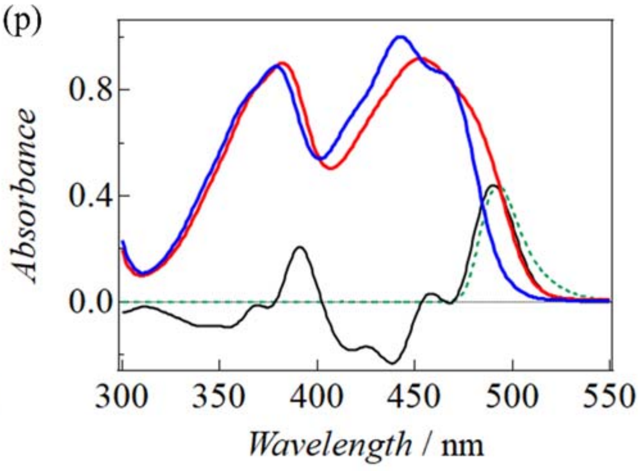
0.8
0.4
0.0
Absorbance
300
350
400
450
500
550
Wavelength / nm

(q)

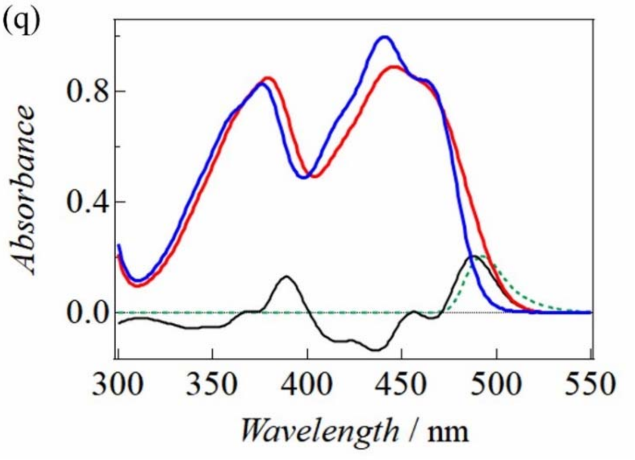
0.8
0.4
0.0
Absorbance
300
350
400
450
500
550
Wavelength / nm

(r)

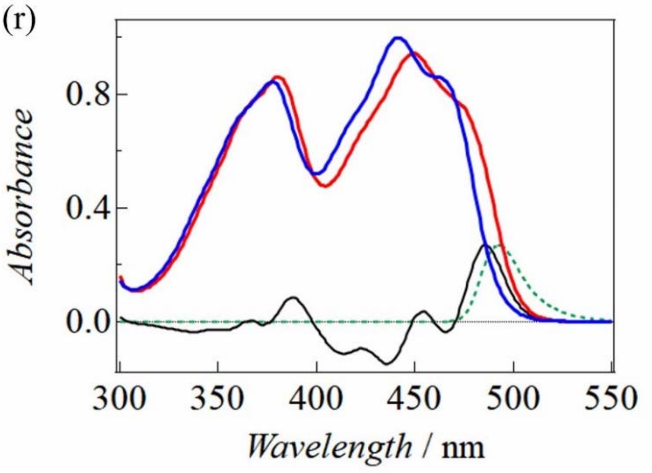
0.8
0.4
0.0
Absorbance
300
350
400
450
500
550
Wavelength / nm

(s)

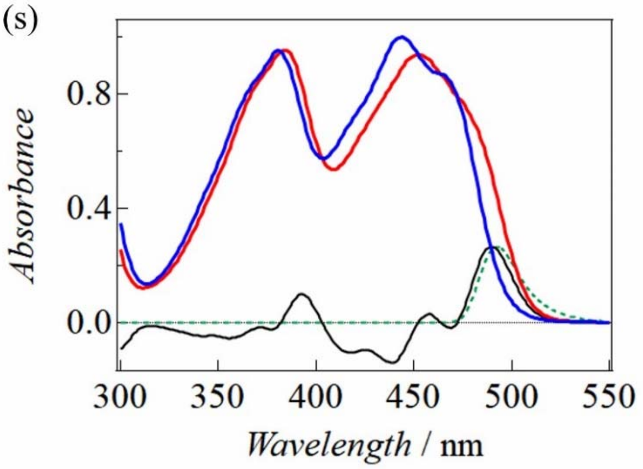
0.8
0.4
0.0
Absorbance
300
350
400
450
500
550
Wavelength / nm

**Fig.S1** The absorption spectra in the dark (blue) and light (red) states for all samples used in this study. In addition, their difference spectra (black) and the profiles of the probe light (green) used in the transient absorption experiments are also shown. The absorption spectra are normalized at a maximum wavelength of around 460 nm in the dark state. (a) $AppA_{5-125}$, (b) $AppA_{1-156}$, (c) $AppA_{1-398}$, (d) OaPAC-BLUF, (e) OaPAC-α3, (f) OaPAC-FL, (g) BlrP1, (h) YcgF, (i) PapB, (j) SyPixD, (k) TePixD, (l) SyTe, (m) TeSy, (n) SyPixD-N31A, (o) SyPixD-N32A, (p) SyPixD-L41V, (q) SyPixD-I66V, (r) SyPixD-W91A, (s) SyPixD-M93A

### SI-3 Definition of $E_{red}$

The amount of the red-shift, $E_{red}$, is defined by the energy difference between the half absorbances of the peaks of the absorption spectra in the light and dark states.

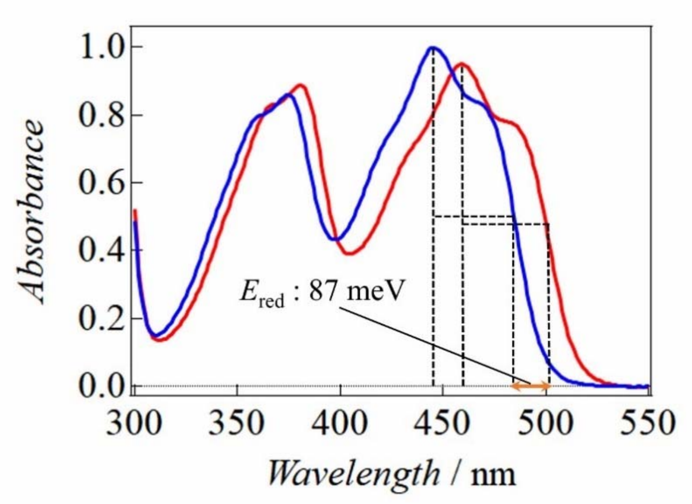


**Fig.S2** An example of $E_{red}$ for $AppA_{5-125}$

### SI-4 Slow rise components of SyPixD at various temperatures

As described in the main text, the slow absorption changes of the BLUF proteins in milliseconds are reproduced by a sum of two exponential functions (two-phase reaction). Initially, we speculated that the two-phase reaction is originated from structural heterogeneities in the BLUF domain. For example, structural heterogeneities were reported for SyPixD, which has a decameric structure with double pentamer rings.[3] The structure determined by the X-ray crystallography showed that the subunit structure of the decamer possesses two conformations of the side chains of W91 ($W_{in}$, $W_{out}$) in the BLUF domain. The occupations of the $W_{in}$ and $W_{out}$ conformations were 1:9 and 2:8 for the monoclinic and tetragonal crystal forms, respectively. It may be possible that these different structures are origin of the different reaction rates. If these heterogeneities are the origin of the two-phase, the amplitude ratio ($A_1/A_2$) should reflect the relative populations of the two conformations. To examine this possibility, the temperature dependence of this ratio is measured. If the two-phase change is originated from the two different conformations and the energies of these structures are different, the relative population of these conformations and $A_1/A_2$ should be temperature dependent.

The transient absorption signals of SyPixD measured at 10-30 °C are shown in Fig.S3. The value of $A_1/A_2$ is almost temperature independent (0.22) within the temperature range (Table SI-1). Although we cannot exclude accidental coincidence of the energies of different conformations, we consider that this possibility is less plausible. In fact, a calculation study based on the crystal structure of SyPixD shows that the energy of $W_{in}$ and $W_{out}$ structures are different.[4] Hence, we believe that the two-phase reaction is originated from two sequential processes of the BLUF protein reaction. This assignment is consistent with the various observations in the main text.

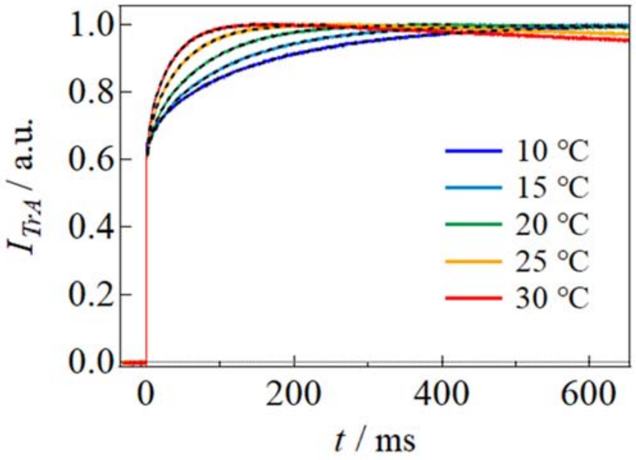


**Fig.S3** Temperature dependence of transient absorption signal of SyPixD. The fitting curves by eq.(1) are shown (broken lines).

Table SI-1

The parameters obtained from the fitting of the transient absorption data of SyPixD, shown in Fig.S3.

| | $A_1$ | $\tau_1$ (ms) | $A_2$ | $\tau_2$ (ms) | $A_1/A_2$ |
|---|---|---|---|---|---|
| 10 °C | 0.068 | 26.8 | 0.30 | 184 | 0.23 |
| 15 °C | 0.067 | 13.5 | 0.31 | 116 | 0.22 |
| 20 °C | 0.071 | 8.09 | 0.32 | 78.6 | 0.22 |
| 25 °C | 0.073 | 5.18 | 0.32 | 48.4 | 0.23 |
| 30 °C | 0.069 | 3.17 | 0.32 | 32.1 | 0.22 |

**SI-5 Transient absorption signals of SyPixD mutants**

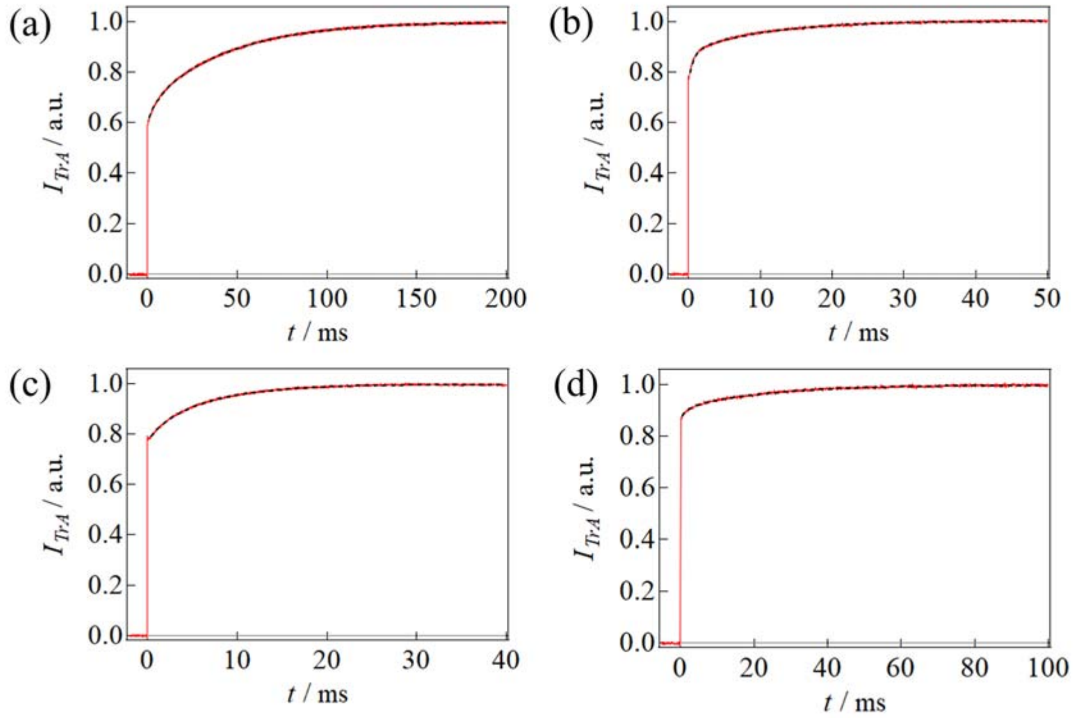


**Fig. S4** Transient absorption signals (red curves) of SyPixD-mutants; (a) N31A (b) L41V (c) I66V (d) M93A. The fitting curves by eq. (1) are shown (broken lines).

### SI-6 Transient absorption signals of W/A mutants of $AppA_{1\text{-}398}$ and OaPAC

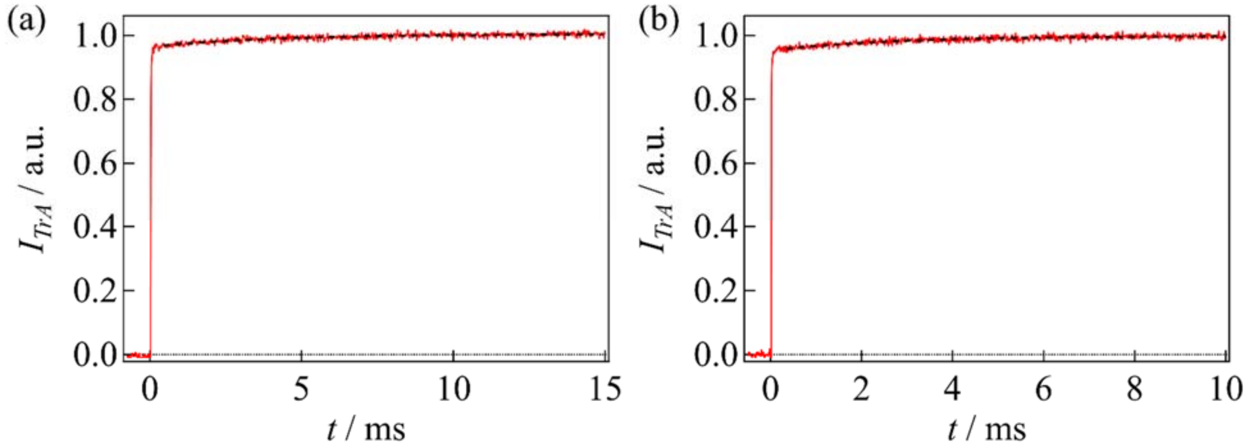


**Fig. S5** Transient absorption signals (red curves) of (a) $AppA_{1\text{-}398}$-W104A mutant, and (b) OaPAC full-length W90A mutant. The fitting curves by eq. (1) are shown (broken lines).

#### References


(1) Winkler, A.; Heintz, U.; Lindner, R.; Reinstein, J.; Shoeman, R. L.; Schlichting, I. A Ternary AppA-PpsR-DNA Complex Mediates Light Regulation of Photosynthesis-Related Gene Expression. *Nat.*

*Struct. Mol. Biol.* **2013**, *20* (7), 859–867.

(2) Shibata, K.; Nakasone, Y.; Terazima, M. Photoreaction of BlrP1: The Role of a Nonlinear Photo-Intensity Sensor. *Phys. Chem. Chem. Phys.* **2018**, *20* (12), 8133–8142.

(3) Yuan, H.; Anderson, S.; Masuda, S.; Dragnea, V.; Moffat, K.; Bauer, C. Crystal Structures of the Synechocystis Photoreceptor Slr1694 Reveal Distinct Structural States Related to Signaling. *Biochemistry* **2006**, *45* (42), 12687–12694.

(4) Goings, J. J.; Reinhardt, C. R.; Hammes-Schiffer, S. Propensity for Proton Relay and Electrostatic Impact of Protein Reorganization in Slr1694 BLUF Photoreceptor. *J. Am. Chem. Soc.* **2018**, *140* (45), 15241–15251.
